## Supplemental files for "Front–rear polarity of intracellular signaling uncovered via giant *Dictyostelium* cells"

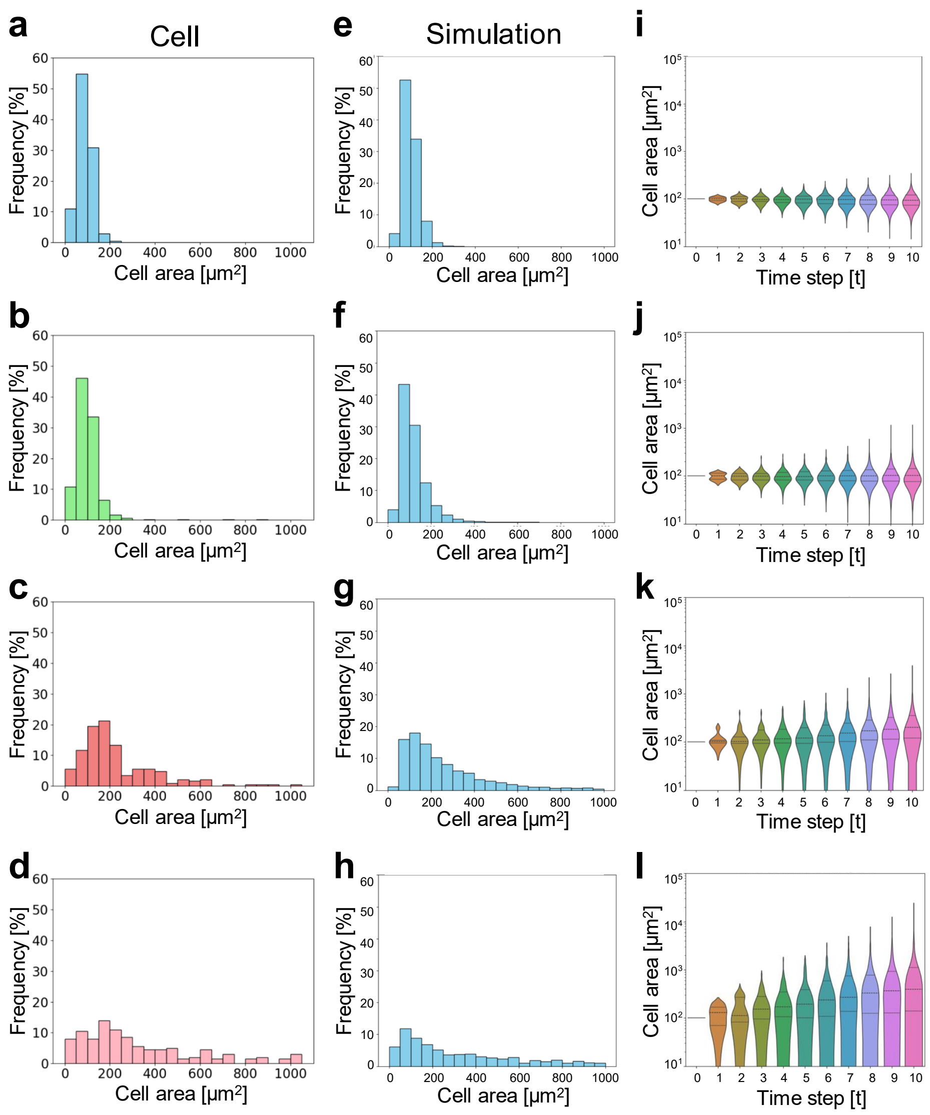


**Extended Data Fig. 1 | Distribution and temporal changes of cell size in simulation and experiment.**

**a–d**, Histograms showing the distribution of cell sizes from experiments. a–d show the same data as presented in Fig. 1a–d, respectively. **e–h**, Simulation results of cell size distributions based on a discrete-time stochastic model are shown. These are similar to the experimental data presented in Fig. 1a–d, respectively. All simulations represent the state after 10 time steps. In each simulation, the initial number of cells $N_0$ was set to 10, and the initial size $S_0$ was 100. The minimum size required for division ($S_{\min}$) was fixed at 50. Panel e shows the case where the probability of cell division $P_{\mathrm{div}}$ was 1.00 and the division ratio $p$ ranged from 0.40 to 0.60. Panel f represents the condition with $P_{\mathrm{div}}$ = 0.95 and the same division ratio range (0.40 to 0.60). Panel g depicts the case with a lower division probability of 0.80, maintaining the same $p$ range. Panel h further decreases $P_{\mathrm{div}}$ to 0.60 and applies a broader division ratio range from 0.20 to 0.80. **i–l**, The progression of cell size distributions over each time step in the simulations shown in panels e–h, respectively.

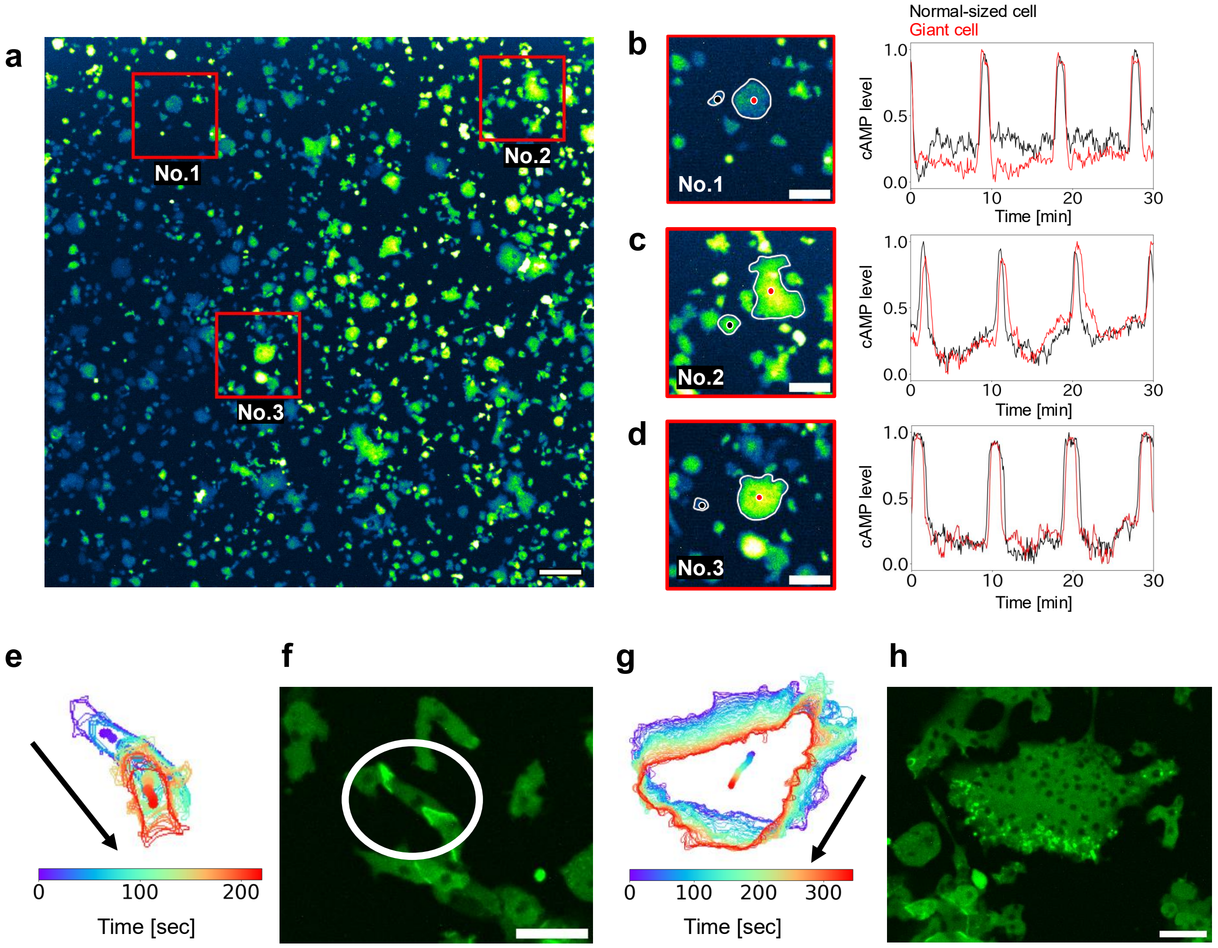


**Extended Data Fig. 2 | Comparison of cell dynamics in cAMP relay between normal-sized and giant cells.**

AX2 cells expressing Flamindo2 were cultured in HL5 medium with 10 μM blebbistatin under shaking conditions for 8 days. After washed and starved in DB, fluorescence imaging was performed using confocal microscopy (10× objective, 5-sec interval). **a**, A fluorescence image showing the entire field of view. Scale bar: 100 μm. Regions where giant and normal-sized cells were adjacent are highlighted in red boxes. **b–d**, cAMP signal dynamics in individual cells (ROIs No.1, 2, and 3). Left panels: Enlarged views of the red boxes shown in (a). Scale bar: 50 μm. Black dots: ROIs in normal-sized cells (diameter: 5.5 μm). Red dots: ROIs in giant cells (diameter: 5.5 μm). Right panels: Plots of normalized cAMP levels over time. Horizontal axis: time [min]; vertical axis: normalized cAMP level. Black lines: normal-sized cells; red lines: giant cells. **e–h**, AX2 cells expressing CRAC-GFP were cultured in HL5 medium with 10 μM blebbistatin for 5 days under shaking conditions. After washed and starved in DB, fluorescence imaging was performed using confocal microscopy (40× objective, 5-sec interval). Centroid trajectories of normal-sized (e) and giant (g) cells. Arrows indicate the direction of migration. Fluorescence images showing CRAC localization in normal-sized (f) and giant (h) cells. Scale bars: 20 μm.

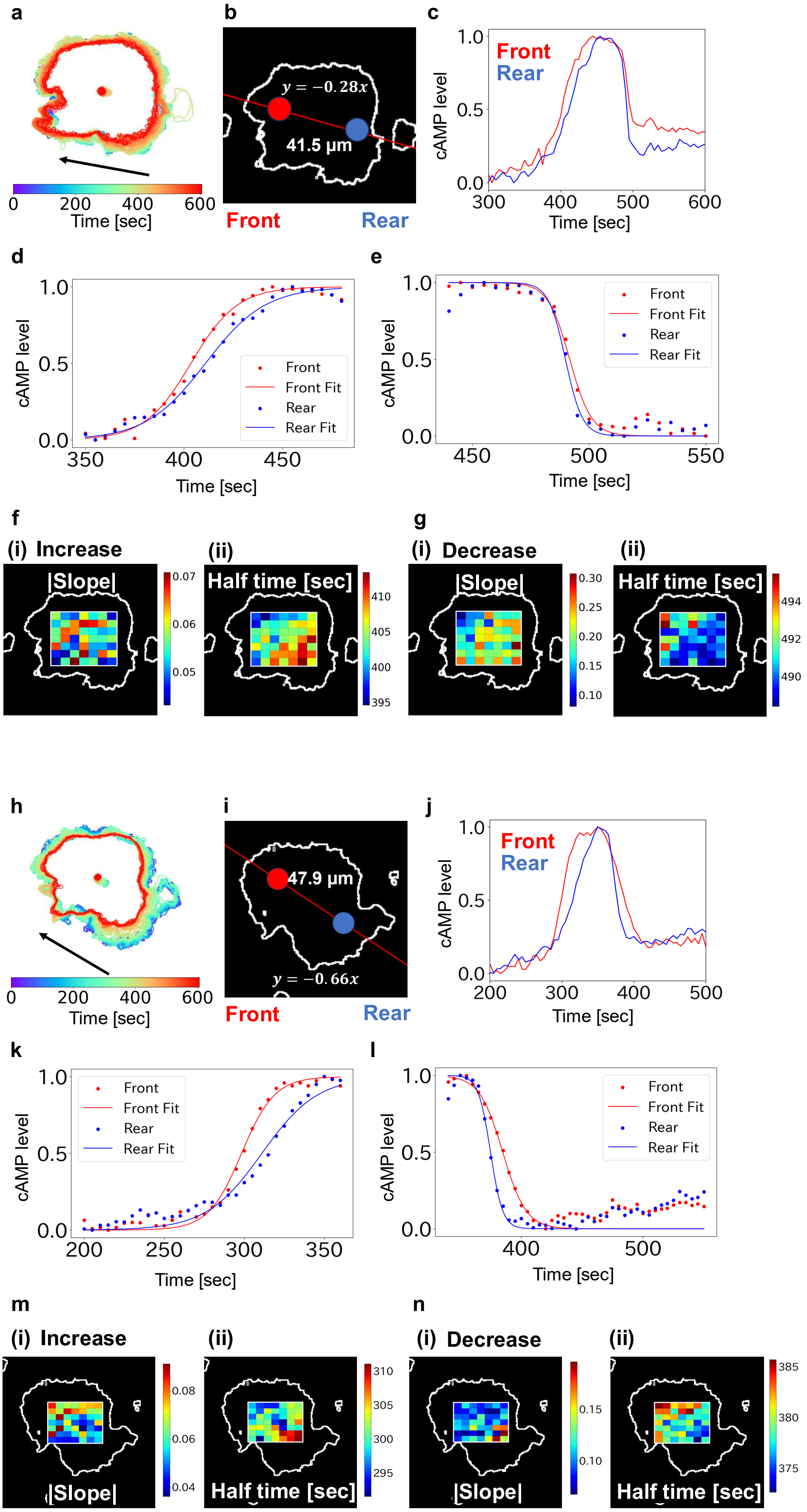


**Extended Data Fig. 3 | Spatiotemporal analysis of cAMP synthesis and decrease.**

AX2 cells expressing Flamindo2 were cultured with 10 μM blebbistatin under shaking conditions for 8 days and then collected at 1×$10^6$ cells/mL. After washed and starved with DB, confocal imaging was then initiated (40× objective, 5-sec intervals). **a–g**, Analysis of a giant cell with a trajectory slope of −0.28. **a**, Cell outline overlaid with centroid trajectory. An arrow indicates the direction of migration. **b**, Analysis regions for the front (red) and rear (blue) were defined based on linear fitting of the trajectory. Regions were located 20 μm from the centroid along the fitted line, with a slope of −0.28. Diameter: 11 μm. **c**, Temporal changes in cAMP levels at the front (red) and rear (blue). Horizontal axis: time [sec]; vertical axis: normalized inverted cAMP intensity. **d**, Sigmoid fitting of cAMP increase. Red: front; blue: rear. **e**, Sigmoid fitting of cAMP decrease using the same equation. **f**, (i) Heatmap of slope parameter during cAMP increase. (ii) Heatmap of half-time parameter during cAMP increase. **g**, (i) Heatmap of slope parameter during cAMP decrease. (ii) Heatmap of half-time parameter during cAMP decrease. **h–n**, Analysis of a different giant cell with a trajectory slope of −0.66. The same experimental procedure was followed as described above. **h**, Cell contour overlaid with centroid trajectory. **i**, Definition of front (red) and rear (blue) regions (slope = −0.66). **j**, Temporal cAMP dynamics at the front and rear. **k**, Sigmoid fitting of the cAMP increase. **l**, Sigmoid fitting of the cAMP decrease. **m**, (i) Heatmap of slope during the cAMP increase. (ii) Heatmap of half-time during the cAMP increase. **n**, (i) Heatmap of slope during the cAMP decrease. (ii) Heatmap of half-time during the cAMP decrease.

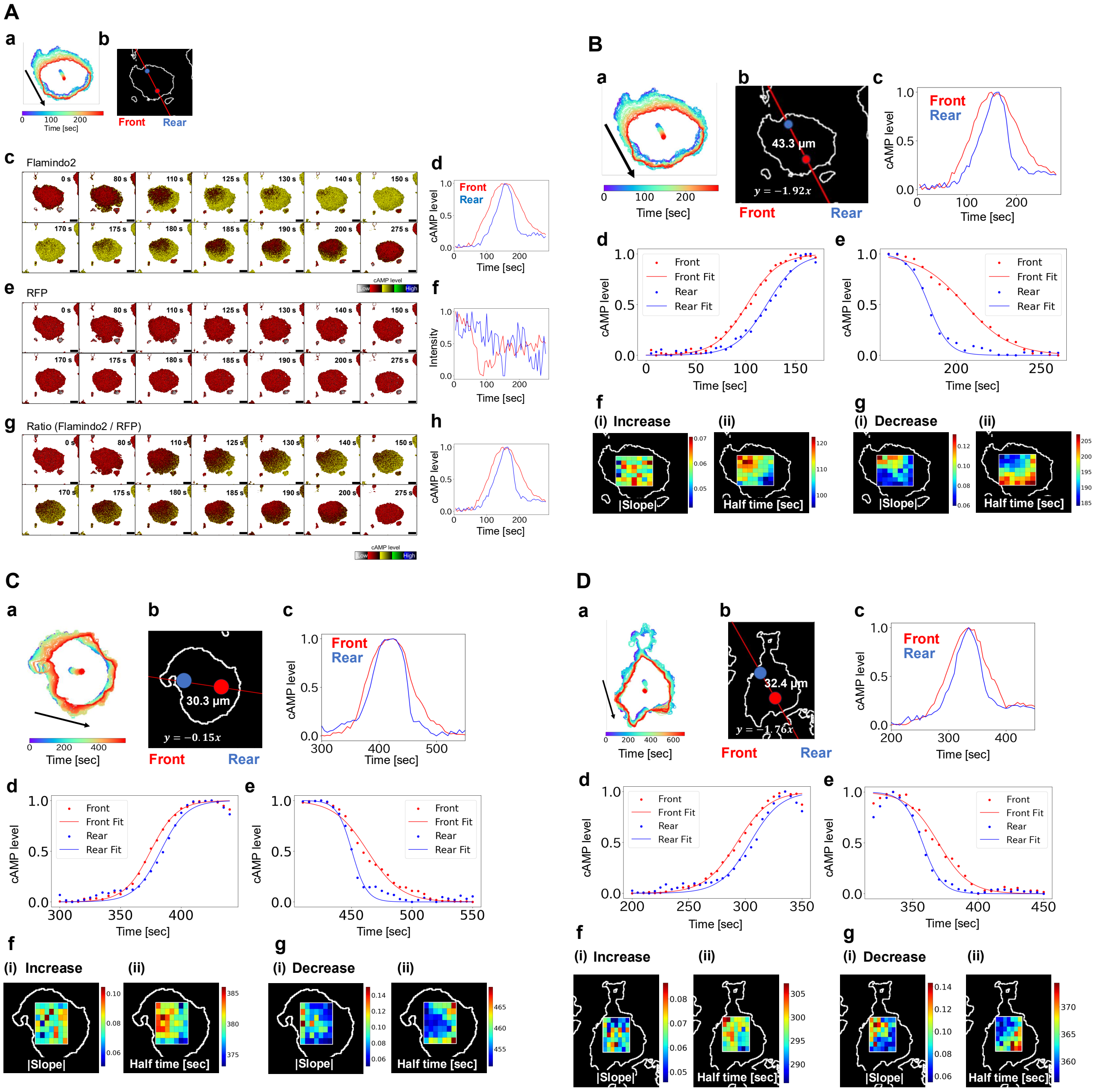


**Extended Data Fig. 4 | Detailed analysis of intracellular cAMP dynamics with Flamindo2-RFP ratiometric imaging.**

AX2 cells expressing Flamindo2-RFP were cultured with 10 μM blebbistatin under shaking conditions. After washed and starved with DB, imaging was performed using confocal microscopy (40× objective, 5-sec intervals).

**A-a**, Cell contour and centroid trajectory of a representative giant cell. An arrow indicates the direction of cell migration. **A-b**, Definition of front and rear regions for analysis. The centroid trajectory obtained in (a) was fitted with a straight line to determine its slope. From the centroid (origin), the front and rear regions were defined by moving 10 μm in the x-direction and ±10 × (−1.92) μm in the y-direction along the fitted slope. Red and blue circles denote the front and rear regions, respectively (diameter: 11 μm). **A-c**, Time-lapse fluorescence images (Flamindo2 channel) of the cell shown in (a). The number in the top right indicates the elapsed time from the start of imaging. Scale bar: 20 μm. **A-d**, Temporal changes in cAMP levels at the front (red) and rear (blue), based on Flamindo2 signal intensity. Data are shown as inverted and normalized fluorescence values. **A-e**, Time-lapse fluorescence images from the RFP channel of the same cell. **A-f**, Fluorescence intensity changes in the RFP channel at the front (red) and rear (blue), representing brightness variation caused by cell movement. **A-g**, RFP/Flamindo2 ratio images to reduce motion artifacts. Representative time-lapse images show corrected intensity at each time point. **A-h**, Corrected cAMP level changes at the front (red) and rear (blue) based on the RFP/Flamindo2 ratio, presented as normalized values. Horizontal axis: time (sec).

Distinct cells were analyzed in panels **B–D**. **B-a**, Cell contour and centroid trajectory of a representative giant cell. Arrows indicate the direction of cell migration. **B-b**, Definition of the front and rear regions based on linear fitting of the centroid trajectory. From the centroid (origin), the front and rear were located by shifting 10 μm in the x-direction and ±10 × (−1.92) μm in the y-direction. Red and blue circles indicate front and rear positions (diameter: 11 μm). **B-c**, Temporal changes in cAMP levels at the front (red) and rear (blue), shown as normalized fluorescence intensity ratios (RFP/Flamindo2). **B-d**, Sigmoid fitting of cAMP increase. Red and blue lines show fitting curves for the front and rear, respectively. **B-e**, Sigmoid fitting of cAMP decrease. **B-f** (i), Heatmap of slope during the increase phase. (ii), Heatmap of half-time during the increase phase. **B-g** (i), Heatmap of slope during the decrease phase. (ii), Heatmap of half-time during the decrease phase.

**C**, Same procedures were applied to another giant cell. **C-a**, Cell contour and centroid trajectory. **C-b**, Definition of front and rear regions based on trajectory (15 μm displacement, slope = −0.15). **C-c**, Temporal changes in cAMP levels at front and rear. **C-d**, Sigmoid fitting of cAMP increase. **C-f**, Sigmoid fitting of cAMP decrease. **C-f** (i–ii), Heatmaps of slope and half-time during the increase phase. **C-g** (i–ii), Heatmaps of slope and half-time during the decrease phase.

**D**, Third cell analysis under the same experimental conditions. **D-a**, Cell trajectory. **D-b**, Front and rear region definitions (8 μm displacement, slope = −1.76). **D-c**, cAMP level dynamics. **D-d**, Sigmoid fitting of increase. **D-e**, Sigmoid fitting of decrease. **D-f** (i–ii), Heatmaps for the increase phase. **D-g** (i–ii), Heatmaps for the decrease phase.

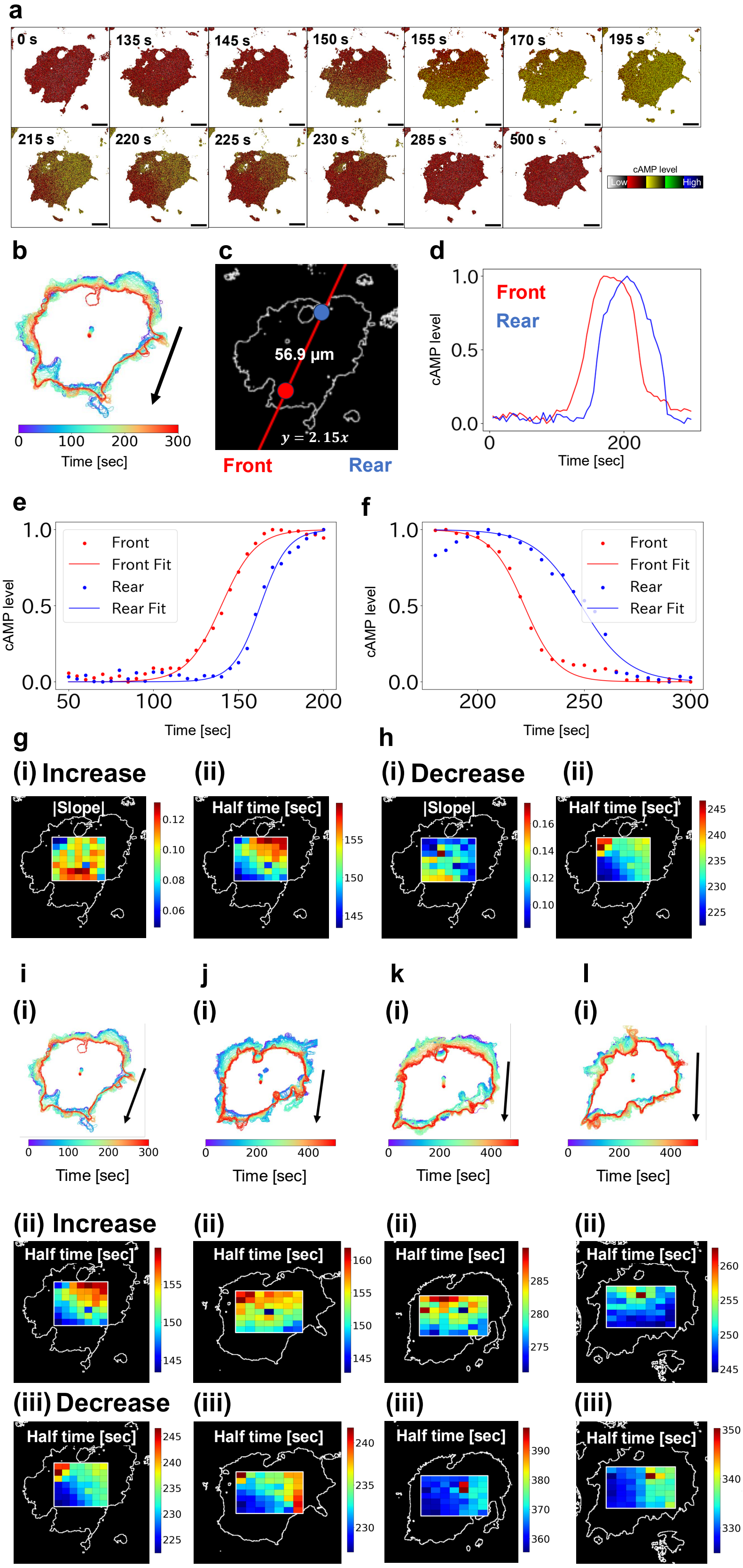


**Extended Data Fig. 5 | Temporal analysis of front-rear polarity in cAMP signaling.**

**a**, Representative RFP/Flamindo2 ratio images of a giant cell extracted from the field of view. Numbers in the top right indicate time elapsed from the start of imaging. Scale bar: 20 μm.

**b**, Overlay of cell contour and centroid trajectory. An arrow indicates the direction of cell migration. **c**, Definition of the front and rear regions for analysis. A linear fit was applied to the centroid trajectory obtained in (b). The front and rear regions were defined by shifting 12 μm in the x-direction and ±12 × 2.15 μm in the y-direction from the centroid, respectively. Red and blue circles indicate the front and rear regions. Diameter: 11 μm. **d**, Temporal changes in cAMP levels at the front (red) and rear (blue) regions. The vertical axis represents normalized fluorescence RFP/Flamindo2 ratio. **e**, Sigmoid fitting of cAMP increase. **f**, Sigmoid fitting of cAMP decrease. **g**, (i) Heatmap of slope during cAMP increase. (ii) Heatmap of half-time during cAMP increase. **h**, (i) Heatmap of slope during cAMP decrease. (ii) Heatmap of half-time (b) during cAMP decrease. **i–l**, (i) Centroid trajectories of the same giant cell during the 1st to 4th signaling events. Arrows indicate the direction of migration. (ii) Corresponding heatmaps of half-time during cAMP increase for each event. (iii) Corresponding heatmaps of half-time during cAMP decrease for each event.

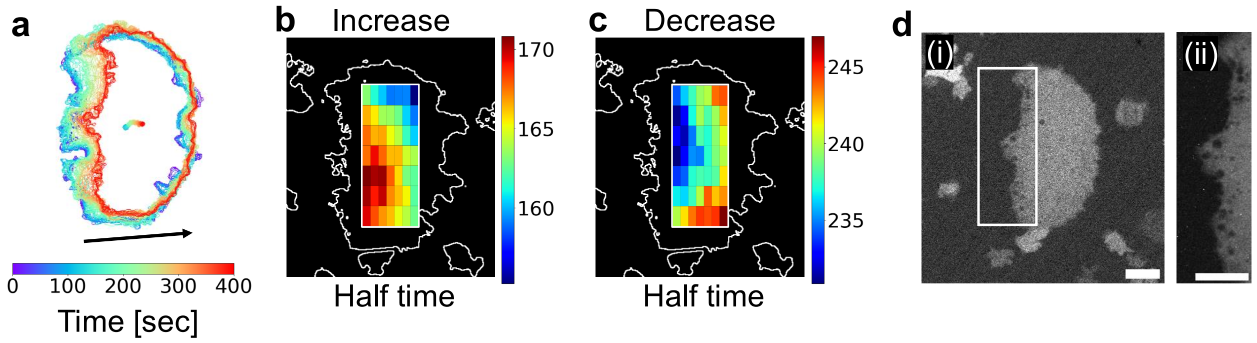


**Extended Data Fig. 6 | Observation of vesicle localization at the rear of giant cells.** **a**, Trajectory of a Flamindo2-RFP giant cell (cultured for 6 days). **b**, Heatmap of half-time during the cAMP increase phase. **c**, Heatmap of half-time during the cAMP decrease phase. **d**, Localization of vesicles at the cell rear. White box indicates magnified area shown in (ii). Scale bar: 20 μm.

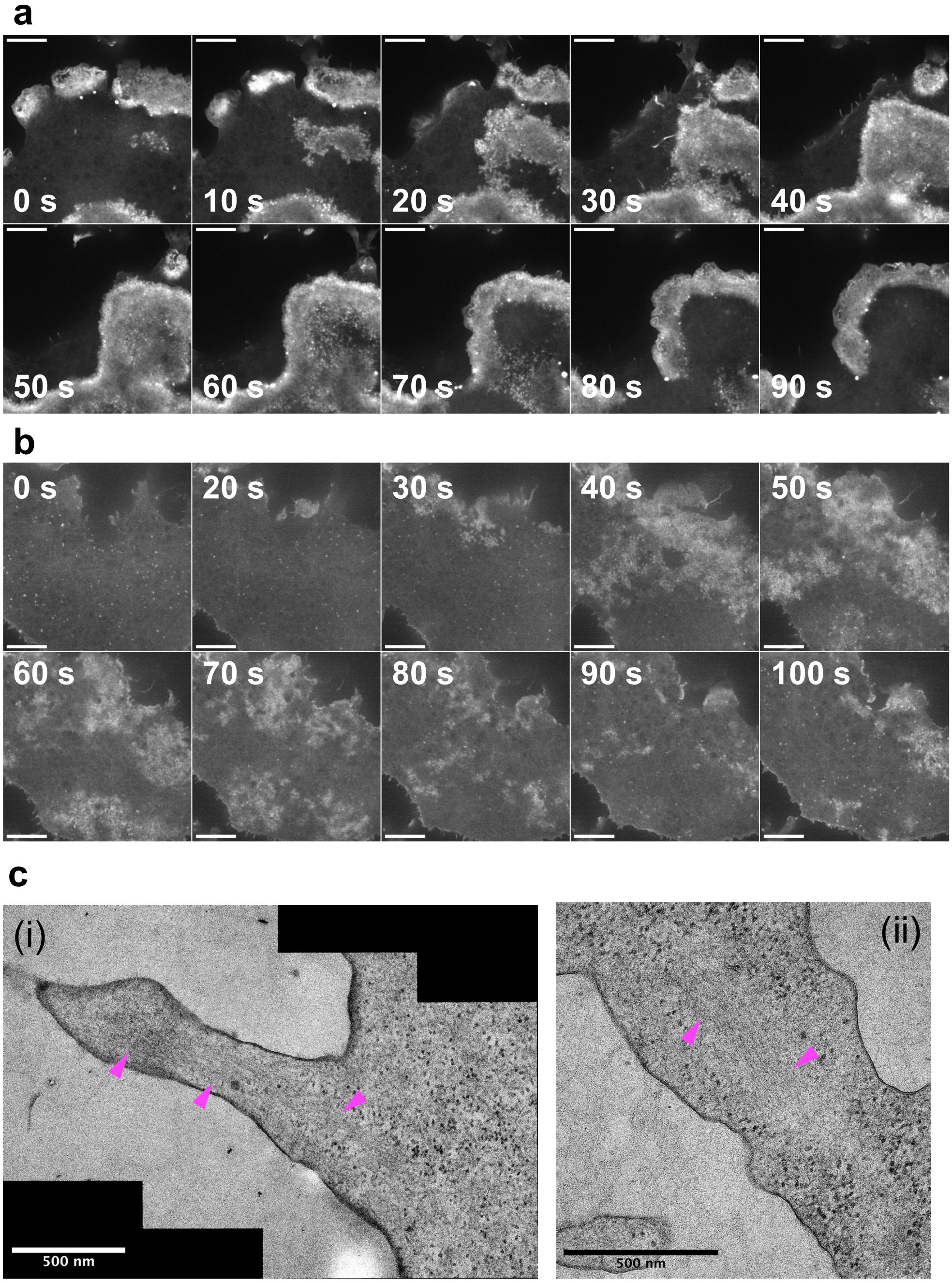


**Extended Data Fig. 7 | Visualization of actin dynamics in giant cells using super-resolution microscopy and electron microscopy.**

AX2 cells expressing Lifeact14-mScarletI were cultured with 10 μM blebbistatin under shaking conditions for 6 days. Imaging was initiated 7.5 hours after starved with DB using confocal microscopy (40× objective, 5-second intervals). **a**, Representative time-lapse Lifeact14-mScarletI fluorescence images of a giant cell. Numbers at the top right indicate time elapsed from the beginning of imaging. Scale bars: 20 μm. **b**, Another representative time-lapse Lifeact14-mScarletI fluorescence images of a giant cell. Scale bars: 20 μm. **c**, Visualization of actin bundle structures (magenta arrowheads) by combining the giant cell preparation method with electron microscopy. Scale bars: 500 nm.

### Legends of Supplementary Videos

**Supplementary Video 1.**

Time-lapse fluorescence imaging of cAMP signal relay was performed in AX2 cells expressing Flamindo2. Giant cells were constructed by blebbistatin treatment and shaking culture. Scale bar: 100 μm. Corresponds to Extended Data Fig. 2a.

**Supplementary Video 2.**

Time-lapse fluorescence imaging of $Ca^{2+}$ oscillations in *Dictyostelium* cells expressing GCaMP6s. Giant cells were constructed by blebbistatin treatment and shaking culture. Scale bar: 100 μm.

**Supplementary Video 3.**

Time-lapse fluorescence imaging of front-to-rear propagation of intracellular cAMP signal in a migrating giant cell expressing Flamindo2. Scale bar: 20 μm. Corresponds to Fig. 2.

**Supplementary Video 4.**

Time-lapse fluorescence imaging of intracellular cAMP signal in a migrating giant cell expressing Flamindo2. Scale bar: 20 μm. Corresponds to Fig. 3h.

**Supplementary Video 5.**

Spatiotemporal variation in cAMP signal initiation and decay in *Dictyostelium* cells expressing Flamindo2-RFP. Scale bar: 20 μm. Corresponds to Extended Data Fig. 5a.

**Supplementary Video 6.**

Micropipette-based cAMP stimulation reveals that signal synthesis initiates near the stimulus. Scale bar: 50 μm. Corresponds to Fig. 4b.

**Supplementary Video 7.**

Biphasic $Ca^{2+}$ dynamics in response to cAMP in a giant cell. GCaMP6s imaging captures dual $Ca^{2+}$ peaks per cAMP cycle. Scale bar: 20 μm. Corresponds to Fig. 6a.

**Supplementary Video 8.**

Spatiotemporal coupling of $Ca^{2+}$ signals and actin wave propagation. Simultaneous imaging of GCaMP6s and Lifeact14-mScarletI shows that actin waves emerge following the reduction of $Ca^{2+}$ levels, indicating inverse coordination between signaling and cytoskeletal dynamics. Scale bar: 20 µm. Corresponds to Fig. 7e.

**Supplementary Video 9.**
Super-resolution imaging of actin structures in a giant *Dictyostelium* cell. SoRA spinning disk confocal microscopy reveals fine actin meshworks in enlarged migrating cells. Scale bar: 10 µm. Corresponds to Extended Data Fig. 7a.

**Supplementary Video 10.**
Super-resolution imaging of actin structures in a giant *Dictyostelium* cell. Scale bar: 10 µm. Corresponds to Extended Data Fig. 7b.
